# ZNF416 is a pivotal transcriptional regulator of fibroblast mechano-activation

**DOI:** 10.1101/2020.07.23.218842

**Authors:** Dakota L. Jones, Jeffrey A. Meridew, Merrick T. Ducharme, Katherine L. Lydon, Kyoung Moo Choi, Nunzia Caporarello, Qi Tan, Patrick A. Link, Ana Maria Diaz Espinosa, Yuning Xiong, Jeong-Heon Lee, Zhenqing Ye, Huihuang Yan, Tamas Ordog, Giovanni Ligresti, Xaralabos Varelas, Daniel J. Tschumperlin

## Abstract

Matrix stiffness is a central regulator of fibroblast function. However, the transcriptional mechanisms linking matrix stiffness to changes in fibroblast phenotype are incompletely understood. Here, we evaluated the effect of matrix stiffness on genome-wide chromatin accessibility in freshly-isolated lung fibroblasts using assay for transposase-accessible chromatin followed by sequencing (ATAC-seq). We found higher matrix stiffness profoundly increased global chromatin accessibility relative to lower matrix stiffness, and these alterations were in close genomic proximity to known pro-fibrotic gene programs. Motif analysis of these regulated genomic loci identified ZNF416 as a putative mediator of fibroblast stiffness responses. Similarly, motif analysis of the promoters of differentially expressed genes observed in freshly sorted fibroblasts from an experimental bleomycin lung fibrosis model also identified ZNF416 as a putative mediator of *in vivo* fibroblast activation. Genome occupancy analysis using chromatin-immunoprecipitation followed by sequencing (ChIP-seq) confirmed that ZNF416 occupies a broad range of genes implicated in fibroblast activation and tissue-fibrosis, with relatively little overlap in genomic occupancy with other mechanoresponsive and pro-fibrotic transcriptional regulators. Using loss and gain of function studies we demonstrated that ZNF416 plays a critical role in fibroblast proliferation, extracellular matrix synthesis and contractile function. Together these observations identify ZNF416 as novel mechano-activated transcriptional regulator of fibroblast biology.

## Introduction

Fibroblasts are tissue-resident mesenchymal cell populations responsible for maintenance and remodeling of the extracellular matrix (ECM). In the case of tissue injury or insult, transient fibroblast activation is critical for proper wound repair^1^. Following injury resolution, activated fibroblasts revert to their quiescent state or undergo apoptosis. Persistent fibroblast activation leads to enhanced ECM deposition and progression of fibro-contractile diseases^2^. In the case of idiopathic pulmonary fibrosis (IPF), sustained fibroblast activation leads to replacement of functional alveolar tissue architecture with scar tissue impairing lung function.^1,3^

Fibroblasts respond to the rigidity of their extracellular surroundings both *in vitro* and *in vivo* such that increased local tissue stiffness acts as an amplification feedback loop driving fibrosis progression^4–10^. For example, fibroblasts in a rigid microenvironment display robust transcriptomic changes^9,11^, accompanied by enhanced ECM deposition and remodeling^9^, amplified proliferation capacity^12^, and increased contractility^12,13^ compared to their counterparts in a compliant microenvironment. Importantly, prior work has shown that primary cells can acquire an epigenetic “mechanical memory” after prolonged culture in rigid mechanical environments ^8,16–18^, suggesting that the use of immortalized or serially-passaged cells could hamper efforts to identify the full regulatory programs that determine fibroblast transcriptional mechanoresponses relevant to *in vivo* processes such as wound healing and fibrosis.

External mechanical stimuli are relayed into intracellular biochemical cascades which ultimately converge onto transcriptional regulators^2^. While YAP/TAZ and MRFT-A are highly studied mechanosensing transcriptional regulators implicated in fibroblast function^2,6,14^ additional transcriptional regulatory mechanisms of mechanosensing likely remain to be identified, and could serve as targets for reversing pathogenic fibroblast activation. The use of epigenomic profiling techniques, such as assay for transposase-accessible chromatin followed by next-generation sequencing (ATAC-seq) has emerged as a method for identifying transcriptional regulators governing epigenetic/transcriptional responses to stimuli. For example, recent ATAC-seq experiments have identified matrix stiffness-dependent chromatin-accessibility changes, specifically in an *in vitro* three-dimensional model of epithelial breast cancer invasion^15^.

Here, we globally assessed the effect of matrix stiffness on chromatin accessibility in freshly isolated mouse lung fibroblasts seeded onto rigid and compliant surfaces. Using *de novo* motif analysis we identified a novel putative mechanoresponsive transcriptional regulator, ZNF416. Analysis of the transcriptional responses in fibroblasts isolated after *in vivo* experimental lung fibrosis in mice were consistent with a potential *in vivo* role for ZNF416 in pathological fibroblast activation. The global DNA occupancy of ZNF416, assessed experimentally by chromatin-immunoprecipitation followed by next-generation sequencing (ChIP-seq), identified interactions of ZNF416 with a wide range of genes critical to pathological fibroblast function. Finally, gain and loss of function studies in lung fibroblasts confirmed a central role for ZNF416 in fibroblast proliferation, contraction, and ECM deposition. Taken together, these results demonstrate the important role of ZNF416 in lung fibroblast function and identify a new transcriptional mechanism by which matrix stiffness regulates cell function.

## Methods

### Primary cell isolation, cell sorting, and cell culture

Primary mouse lung fibroblasts were isolated as previously described by Caporarello, et al^19^ under a protocol approved by the Mayo Clinic Institutional Animal Care and Use Committee (IACUC). *Col1α1*-GFP transgenic mice generated as previously described^20^ were kindly provided by Dr. Derek Radisky. Briefly, 6-8 week old *Col1α1*-GFP mice were anesthetized with ketamine/xylazine solution (100mg/kg and 10mg/kg, respectively) injected intraperitoneally. The left ventricle of the heart was perfused with ice cold PBS (Thermo Fisher Scientific, Waltham, MA, USA) to remove blood content from the lung. Lungs were then immediately harvested and minced in 10-centimeter petri dishes and then incubated in digestive solution (DMEM medium, 0.2 mg/mL Liberase DL, 100 U/mL DNase I). Samples were digested at 37 °C for 45 minutes. Digestive solution was inactivated with 1x DMEM (Thermo Fisher Scientific, Waltham, MA, USA) containing 10% FBS (Thermo Fisher Scientific, Waltham, MA, USA). Cell and tissue suspension was put through a 40 μm filter and centrifuged. Cell pellet was resuspended in red blood cell lysis buffer (BioLegend, San Diego, CA, USA) for 90 seconds and then diluted in 3x volume of PBS. Cells were centrifuged and resuspended in 200 μL of FACS buffer (1% BSA, 0.5 mM EDTA, pH 7.4 in PBS). The single cell suspension was then incubated with anti-CD45:PerCp-Cy5.5 (BioLegend, San Diego, CA, USA) (1:200), anti-CD31-PE (BioLegend, San Diego, CA, USA) (1:200), anti-EpCAM-APC (BioLegend, San Diego, CA, USA) (1:200), and DAPI (Sigma Aldrich, St. Louis, MO, USA) (1:1000) antibodies for 30 minutes on ice. All antibody information is provided in Table S1.

Samples were subjected to fluorescence-activated cell sorting (FACS) using a BD FACS Aria II (BD Biosciences, San Jose, CA, USA). To isolate the CD45-(hematopoietic), EpCAM-(epithelial), CD31-(endothelial), GFP+ (collagen I-expressing) population the following isolation strategy was used: debris exclusion (FSC-A by SSC-A), doublet exclusion (SSC-W by SSC-H and FSC-W by FSC-H), dead cell exclusion (DAPI by anti-CD31-PE), CD45+ cell exclusion (anti-CD45-PerCP-Cy5.5 by *Col1a1-* GFP), EpCAM, and CD31+ cells exclusion (anti-CD325-APC by anti-CD31-PE), and isolation of GFP+ cells (APC by GFP). A schematic of sorting strategy is provided in Figure S1. Primary mouse fibroblasts were cultured in DMEM (Thermo Fisher Scientific, Waltham, MA, USA) supplemented with 10% FBS (Thermo Fisher Scientific, Waltham, MA, USA) and Anti-Anti (Thermo Fisher Scientific, Waltham, MA, USA) unless otherwise stated.

Primary human lung fibroblasts were isolated by explant culture from the lungs of subjects who underwent lung transplantation and were kindly provided by Peter Bitterman at the University of Minnesota under a protocol approved by the University of Minnesota Institutional Review Board. Primary human lung fibroblasts were maintained in DMEM (Thermo Fisher Scientific, Waltham, MA, USA) supplemented with 10% FBS (Thermo Fisher Scientific, Waltham, MA, USA) and Anti-Anti (Thermo Fisher Scientific, Waltham, MA, USA).

### ATAC-seq and analysis

FACS sorted mouse GFP+ lung fibroblasts were seeded onto 0.2 kPa PDMS, 32 kPa PDMS (Advanced Biomatrix, San Diego, CA, USA), or tissue-culture plastic, all identically coated with collagen I (Advanced Biomatrix, San Diego, CA, USA), and maintained for 8 days. Fibroblasts were then trypsinized and counted. 50,000 cells were subjected to Omni ATAC-seq following the published protocol^21^. The size of library DNA was determined from the amplified and purified library by a Fragment Analyzer (Advanced Analytical Technologies; AATI; Ankeny, IA), and the enrichment of accessible regions was determined by the fold difference between positive and negative genomic loci using real-time PCR. The following primer sequences were used: accessibility-positive control locus: AT-P7-F: 5’- GGCTTATCCGGAGCGGAAAT -3′, AT-P7-R: 5’- GGCTGGAACAGGTTGTGTTG -3′. Accessibility-negative control locus: AT-P13-F: 5’ -TCCCCTTTACTGTTTTCCTCTAC -3′, AT-P13-R: 5’ -GGATTGATGAGGAAACAGCCTC -3′. The libraries were sequenced to 51 base pairs from both ends on an Illumina HiSeq 4000 instrument (Illumina, San Diego, CA, USA).

Paired-end reads were mapped to the mm10 genome using BWA^22^. Sam files were converted to Bam files using picard SortSam (https://broadinstitute.github.io/picard/command-line-overview.html#SortSam) and sorted by chromosomal coordinates. PCR duplicates were removed using picard MarkDuplicates (https://broadinstitute.github.io/picard/command-line-overview.html#MarkDuplicates). Pairs of reads with one or both reads uniquely mapped to chr1-22, chrX and chrY were retained. Processed bam files were used to call peaks using MACS2 with the following options “--keep-dup all -q 0.01 -no model”^23^. To identify differentially accessible genomic loci, DiffBind was used with the processed bam files and peak files obtained from MACS2 using an FDR threshold of 0.05^24^. Differentially accessible loci were then annotated to their closest transcriptional start site (TSS) using Homer^25^. Gene ontology was performed using Panther on the genes assigned to differential chromatin accessibility sites. Motif analysis was completed using the findMotifsGenome.pl command within Homer.

### *In vivo* model of experimental-lung fibrosis and motif analysis

To evaluate motif enrichment from fibroblasts activated *in vivo*, we analyzed an unpublished data set from previously completed experiments. These animal experiments were carried out under protocols approved by the Mayo Clinic Institutional Animal Care and Use Committee (IACUC). Two-month old *Col1a1*-GFP male mice were administered bleomycin (1.2U/kg) or PBS intratracheally as described previously^26^. Mice were sacrificed 14 days after bleomycin exposure to allow lung fibrosis to manifest. Mice were anaesthetized, euthanized, and lung GFP+ fibroblasts were FACS sorted as detailed above. GFP+ lung fibroblasts were sorted into RLT lysis buffer and RNA was isolated as detailed below and submitted for sequencing. RNA sequencing data was analyzed as described previously^27^. Briefly, reads were aligned to the mm10 build of the mouse reference genome using STAR with default parameter setting^28^. FeatureCounts was used to generate raw counts as well as normalized RPKM (Reads Per Kilobase of exon per Million mapped reads)^29^. To identify differentially expressed genes in bleomycin-treated mice compared to sham we used an FDR cut-off of 0.05 and a log_2_ fold-change of +/−1. The list of 4,124 genes was submitted to Homer for *de novo* motif enrichment analysis in the promoter region (+/−1kb from TSS). Motifs corresponding to transcription factors not expressed in our fibroblasts (RPKM < 0.01) were discarded from our analysis.

### Microscopy and image analysis

Mouse lung GFP+ fibroblasts were fixed with 4% PFA (Sigma Aldrich, St. Louis, MO, USA) for 15 minutes followed by permeabilization with 0.1% Triton X-100 (Sigma Aldrich, St. Louis, MO, USA) in PBS (Thermo Fisher Scientific, Waltham, MA, USA) for 10 minutes. Samples were then blocked in 5% BSA (Sigma Aldrich, St. Louis, MO, USA) in PBS supplemented with 0.1% Tween-20 (Sigma Aldrich, St. Louis, MO, USA) for 1 hour. Samples were then placed in primary Lamin A/C antibody (Cell Signaling Technology, Danvers, MA, USA) diluted in blocking buffer and were incubated overnight at 4 °C. The following morning, samples were then washed with PBS 3x and then placed in species-specific RFP secondary antibody (Thermo Fisher Scientific, Waltham, MA, USA) for 1 hour. RFP phalloidin (Thermo Fisher Scientific, Waltham, MA, USA) was used to quantify cell area and was done according to manufacturer’s protocol. Samples were then washed with PBS 3x, and Prolong Gold Anti-fade mounting media (Thermo Fisher Scientific, Waltham, MA, USA) was placed on each sample, followed by a small glass coverslip. Samples were then imaged using a z-stack algorithm on a confocal microscope (Zeiss LSM780, Zeiss, Oberkochen, Germany). Z-stack images (0.2 μm per stack) were then reconstructed into 3-dimensional images using Imaris (Bitplane, Zürich, Switzerland) from which nuclear volume was extracted. To measure cell area, confocal RFP phalloidin images were quantified in ImageJ. To measure Ki67 nuclear intensity, nuclei were outlined using DAPI. Fluorescence intensity was then measured within each outlined nuclei. Data are reported as average fluorescence intensity with nuclei subtracted by background. Background is defined as an empty well treated with primary and secondary antibodies.

### RNAi knockdown and lentiviral-mediated overexpression of ZNF416

RNA interference was completed with SMARTpool: ON-TARGETplus siRNA’s (Dharmacon, Lafayette, CO, USA) specific for ZNF416 or scramble non-targeting siRNA as a control. To confirm on target efficacy of the siRNA pool, we tested individual siRNA’s (n=4) and measured ZNF416 transcript levels (Fig. S2). Primary human lung fibroblasts were transfected in 10% FBS in DMEM for 72 hours after which they were used for experiments.

Lentiviral expression plasmids for ZNF416 were purchased from Origene, Inc (Origene, Rockville, MD, USA). Briefly, we used three plasmids: (i) TRC2 pLKO.5-puro empty vector control plasmid (Sigma Aldrich, St. Louis, MO, USA) (ii) ZNF416-GFP (Origene, Inc), and (iii) ZNF416-FLAG (Origene, Inc). Using Lipofectamine 3000, HEK293T cells were transfected with appropriate expression vector along with psPAX (Addgene 12260) and pMD2.G (Addgene 12259) in Opti-MEM (Thermo Fisher Scientific, Waltham, MA, USA) in 5% FBS with 1mM sodium pyruvate (Thermo Fisher Scientific, Waltham, MA, USA) with no antibiotics. Viral media was collected 24 and 48 hours after transfection. To remove potential HEK293T contamination, viral media was spun down at 1,250 rpm for 10 minutes at 4 °C and then passed through a 40 μm PES filter. Viral media with polybrene (10 μg/mL) (Thermo Fisher Scientific, Waltham, MA, USA) was then added to primary human lung fibroblasts and IMR90s, a commercially-available human lung fibroblast cell line. The next day, fibroblasts were washed with PBS and placed in 10% FBS in DMEM with Anti-Anti. Three days after infection, fibroblasts were subjected to puromycin selection (1 μg/mL) (Sigma Aldrich, St. Louis, MO, USA) for 7 days. Following puromycin selection, surviving cells were then validated for expression of ZNF416-GFP and ZNF416-FLAG by western blot (Fig. S3).

### RNA isolation

mRNA was isolated using RNeasy mini kit (Qiagen, Valencia, CA, USA); concentration of mRNA was quantified using a Nanodrop spectrophotometer. cDNA was synthesized using the SuperScript VILO kit (Thermo Fisher Scientific, Waltham, MA, USA). qRT-PCR was completed using FastStart Essential DNA Green Master (Roche Diagnostics, Mannheim, Germany) and analyzed using a LightCycler 96 (Roche Diagnostics, Mannheim, Germany). Primers used for qRT-PCR are listed in Table S2.

### Protein lysis and western blotting

For protein analysis, cells were lysed with RIPA buffer (Thermo Fisher Scientific, Waltham, MA, USA) supplemented with Halt™ protease & phosphatase inhibitor (Thermo Fisher Scientific, Waltham, MA, USA). Protein concentration was quantified with the Pierce BCA Protein Assay kit (Thermo Fisher Scientific, Waltham, MA, USA). Protein samples were loaded onto 4-15% gradient SDS-PAGE gels (Bio-Rad, Hercules, CA, USA), transferred to 0.2 μm pore-size PVDF membranes, and blocked for 1 hour in blocking buffer (5% milk, 0.1% Tween-20 in 1x TBS (Bio-Rad, Hercules, CA, USA)). Blots were then placed into primary antibodies overnight at 4 °C. The following morning, blots were then incubated with species-specific secondary HRP-conjugated antibody for 1 hour at room temperature. All antibody information is located in Table S1. Protein bands were observed using Super Signal West Pico Plus (Thermo Fisher Scientific, Waltham, MA, USA) and images were acquired using a ChemiDoc Imaging System (Bio-Rad, Hercules, CA, USA).

### Collagen gel compaction assays

Cell-embedded collagen micro tissues were generated as previously described^30,31^. Briefly, fibroblasts (4 × 10^6^ cells/mL) were diluted in rat-tail Collagen type-I (6mg/mL) (Corning, Corning, NY, USA). Polystyrene beads (1:50) (Thermo Fisher Scientific, Waltham, MA, USA) 1μm in diameter were added to the solution to visualize the droplet. Emulsions were formed by flow-focusing the collagen/cell solution and oil (fluorocarbon oil FC-40 (Sigma Aldrich, St. Louis, MO, USA)) with 2% FluoroSurfactant (RAN Biotechnologies, Beverly, MA, USA) into a microfluidic PDMS device. Droplet formations were performed at 4°C while the droplets were recovered at 37°C. The fibroblast-containing droplets were incubated at 37°C in medium containing either DMSO (Fisher Scientific, Waltham, MA, USA) or 2ng/mL TGFβ (PeproTech, Rocky Hill, NJ, USA). To quantify the effect of knockdown of ZNF416 on fibroblast compaction, primary human lung fibroblasts were treated with siRNA for 48 hours prior to embedding into collagen microtissues.

### ECM deposition

Fibroblasts were plated in 96-well plates (10,000 fibroblasts/well) for 3 days in 2% FBS in DMEM supplemented with ascorbic acid (50 μg/mL) (Fisher Scientific, Waltham, MA, USA). Fibroblasts and matrix were then fixed and immunostained as detailed above. Briefly, we used Collagen I and Fibronectin antibodies to probe for ECM deposition. Antibody information can be found in Table S1. Species specific secondary antibodies were used as detailed above. Images were acquired using a Cytation 5 microscope (BioTek, Winooski, VT, USA) using a 4x objective. ECM deposition was quantified by RFP image intensity divided by cell number, calculated by DAPI nuclear stain.

### Chromatin-immunopreciptation (ChIP-seq) and analysis

ZNF416-FLAG expressing IMR90 cells were subjected to ChIP-seq for FLAG as previously described^32^. Briefly, IMR90 cells (n=2 biological replicates) stably expressing ZNF416-FLAG were cross-linked with 1% formaldehyde (Sigma Aldrich, St. Louis, MO, USA) for 10 minutes, followed by quenching with 125 mM glycine (Sigma Aldrich, St. Louis, MO, USA) for 5 minutes at room temperature. Fixed cells were washed with 1X TBS (Thermo Fisher Scientific, Waltham, MA, USA). IMR90 cells were then resuspended in cell lysis buffer (10 mM Tris-HCl, pH7.5, 10 mM NaCl, 0.5% NP-40) and incubated on ice for 10 minutes. The lysates were washed with MNase digestion buffer (20 mM Tris-HCl, pH7.5, 15 mM NaCl, 60 mM KCl, 1 mM CaCl_2_) and incubated for 20 minutes at 37 °C in the presence of MNase. After adding the same volume of sonication buffer (100 mM Tris-HCl, pH8.1, 20 mM EDTA, 200 mM NaCl, 2% Triton X-100, 0.2% Sodium deoxycholate), the lysate was sonicated for 10 minutes (30 sec-on / 30 sec-off) in a Bioruptor instrument (Diagenode, Denville, NJ, USA) and then centrifuged at 15,000 rpm for 10 minutes. The cleared supernatant equivalent to about 15 million cells was incubated with 30 μl of prewashed anti-FLAG M2 magnetic beads (Sigma, Catalogue #: M8823) (Sigma Aldrich, St. Louis, MO, USA) on a rocker overnight. The beads were extensively washed with ChIP buffer, high salt buffer, LiCl_2_ buffer, and TE buffer. Bound chromatin was eluted and reverse-crosslinked at 65 °C overnight. DNA was treated with RNase A and Proteinase K (Qiagen, Valencia, CA, USA) and then purified using a Min-Elute PCR purification kit (Qiagen, Valencia, CA, USA). Sequencing libraries were prepared from 5 - 10 ng of ChIP and input DNA with the ThruPLEX® DNA-seq Kit V2 (Rubicon Genomics, Ann Arbor, MI) and were sequenced to 51 base pairs from both ends using an Illumina HiSeq 4000 instrument (Illumina, San Diego, CA, USA).

Paired-end fastq files were aligned to the hg19 build of the human genome using Bowtie2 using the default settings^33^. SAM files were converted to BAM and were sorted by chromosomal-coordinates using Picard SortSam. Duplicates were removed using Picard MarkDuplicates. BAM files were then used to call peaks using MACS2 using default settings with a q threshold of 0.05^23^. To generate BigWig files, deepTools BamCoverage was used with default settings and a bin-size of 10 base pairs^34^. Integrative Genome Viewer (IGV) was used to visualize BigWig files on the hg19 genome track. Homer was used to annotate ZNF416 binding sites to the nearest TSS as described above. Gene ontology of ZNF416 target genes was completed as described above.

To test whether ZNF416-FLAG binding locations are co-occupied by the mechanoresponsive or profibrotic transcription factors YAP, SMAD3, or SRF, or H3K27-acetylation (H3K27ac), an active enhancer mark indicating gene activation, publicly-available ChIP-seq data were downloaded (GSE61852 – YAP^35^, GSE38103 – SMAD3^36^, GSE61475 – SRF^37^, GSE16256 – H3K27ac^38^). Homer was used to identify co-occupancy using the mergePeaks command within Homer. Regions were considered to be co-occupied if the peak distance between two marks was 300 bp or less.

### Statistical analysis and figure preparation

GraphPad Prism 8.0 (GraphPad Software, La Jolla, CA, USA) was used for generating Figures 1-5 and statistical analyses. Data are represented as mean +/−standard error. Statistical analysis was done using either an un-paired two-tailed t-test when directly comparing two groups or one-way ANOVA with Tukey multiple-comparison adjustment when comparing 3 or more groups.

**Figure 1:**
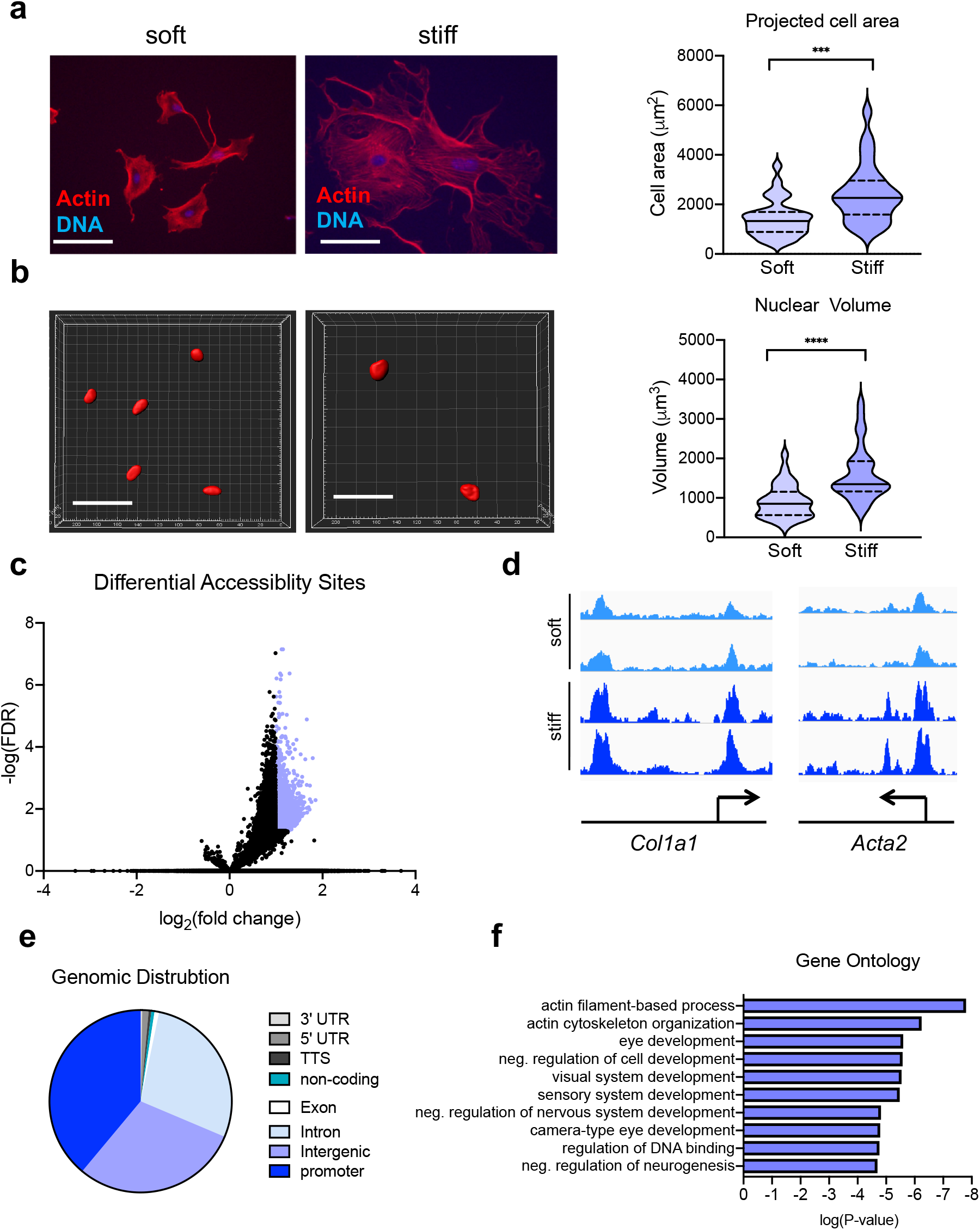
Matrix stiffness facilitates increased chromatin accessibility. (a) 20x fluorescent images of FACS sorted *Col1a1*-GFP mouse lung fibroblasts accompanied with quantification of cell area on soft vs stiff matrices (soft: n=32, stiff: n=27). Solid lines in violin plots represent median. Dotted lines in violin plots represent upper and lower quartiles. (b) Three-dimensional nuclei reconstructed in Imaris accompanied with quantification of nuclear volume on soft vs stiff matrices (soft: n=32, stiff: n=30). Solid lines in violin plots represent median. Dotted lines in violin plots represent upper and lower quartiles. (c) Volcano plot of differential accessibility sites of fibroblasts on soft vs stiff matrices. Each dot represents a single accessibility site. Significantly different accessibility sites colored in blue. (d) Genomic views of differential chromatin accessibility sites in close proximity to known pro-fibrotic genes. (e) Genomic distribution of differential accessibility sites annotated to their nearest TSS. (f) Ontology analysis of genes annotated to all differential accessibility sites. Scale bars represent 50 μm. ***P< 0.001, ****P<0.0001, evaluated by unpaired students t-test.

## Results

### Matrix stiffness increases chromatin accessibility

To address the role of matrix rigidity in shaping the chromatin accessibility landscape of “naïve” fibroblasts, we used a *Col1α1*-GFP reporter mouse to freshly isolate the matrix-producing fibroblasts of the lung (CD45-CD326-CD31-GFP+) using FACS (Fig. S1). These freshly-isolated naïve fibroblasts were placed on identically collagen I coated soft (0.2 kPa PDMS) or stiff (tissue-culture plastic) substrates for 8 days. Naïve fibroblasts were responsive to the rigidity of their substrate as shown by their change in total cell area (Fig. 1a) and nuclear volume (Fig. 1b). To address whether matrix rigidity alters chromatin accessibility of naïve lung fibroblasts, we used ATAC-seq, a fast and efficient next-generation sequencing method to identify active regulatory elements (enhancers and promoters) genome-wide. Using an FDR ≤ 0.05 and a log_2_ fold-change of +/−1, we found that stiff matrices increased accessibility at 2,114 genomic loci compared to soft matrices (Fig. 1c, Table S3). Stiff matrices did not promote any reduction in chromatin accessibility compared to soft matrices (Fig. 1c, Table S3). Analyzing ATACseq data generated in cells cultured on 32 kPa PDMS substrates revealed no significant differences relative to cells cultured on tissue culture plastic, indicating that the latter is a substrate sufficient to serve as our primary model of stiff tissue matrices (Table S4).

Using Homer, we annotated these 2,114 genomic loci to the nearest transcriptional start site. This analysis revealed accessibility changes in close proximity to fibroblast activation genes such as *Col1a1* and *Acta2* (Fig. 1d). The genomic distribution of these differentially accessible sites (DAS) revealed promoter regions as the dominant regulatory regions responding to matrix stiffness (38.9% of DAS), followed by intergenic/enhancer regions (29.7% of DAS), and introns (28% of DAS) (Fig. 1e). Using Panther, we then conducted Gene Ontology analysis of the annotated DAS. We found enrichment of genes involved in “actin filament-based process” and “actin cytoskeleton organization”, consistent with matrix stiffness programmatically engaging fibroblast mechanoresponses through chromatin accessibility changes of these gene programs (Fig. 1f).

### Motif enrichment analysis identifies ZNF416 as a putative regulator of fibroblast activation *in vitro* and *in vivo*

Transcription factors are a class of proteins that have the capacity to bind to specific DNA sequences, or motifs, and then recruit epigenetic machinery to modulate chromatin accessibility at defined genomic loci regulating gene expression. To identify candidate transcription factors associated with matrix stiffness directed changes in chromatin accessibility, we performed *de novo* motif analysis on the 2,114 matrix stiffness DAS using Homer (Fig. 2a). Ranked by ascending P-value, the top predicted transcription factor was Nuclear Factor, Erythroid 2 Like 2 (Nrf2). Interestingly, Nrf2 has been shown to translocate from the cytoplasm to the nucleus in endothelial cells under uniform shear stress *in vitro* demonstrating its role in cell mechanoresponsiveness^39^. Second was FOS Like 1 (Fosl1 or Fra1), a member of the activating-protein 1 complex (AP-1). This transcriptional complex has been implicated in the progression of a variety of experimental tissue-fibrosis models (e.g. lung, kidney, heart)^40^. Third was Zinc Finger Protein 416 (ZNF416) whose function has not yet been defined in the literature. Fourth was SP1, a multifunctional transcription factor whose role in three-dimensional matrix stiffness-induced breast cancer invasion has recently been described^15^. Fifth was TEA Domain Transcription Factor 4 (TEAD4) which mediates transcriptional activity of YAP/TAZ^41,42^, major mechanotransducers that contribute to fibroblast activation and fibrogenesis *in vitro* and *in vivo*^6,7,10,43,44^. Sixth was TAL BHLH Transcription Factor 1 (TAL1) which is considered a driver of T-cell acute lymphoblastic leukemia^45^ and whose function within the context of mechanotransduction has not yet been studied. A complete list of motifs identified in this analysis is provided in Supplemental Table 5.

**Figure 2:**
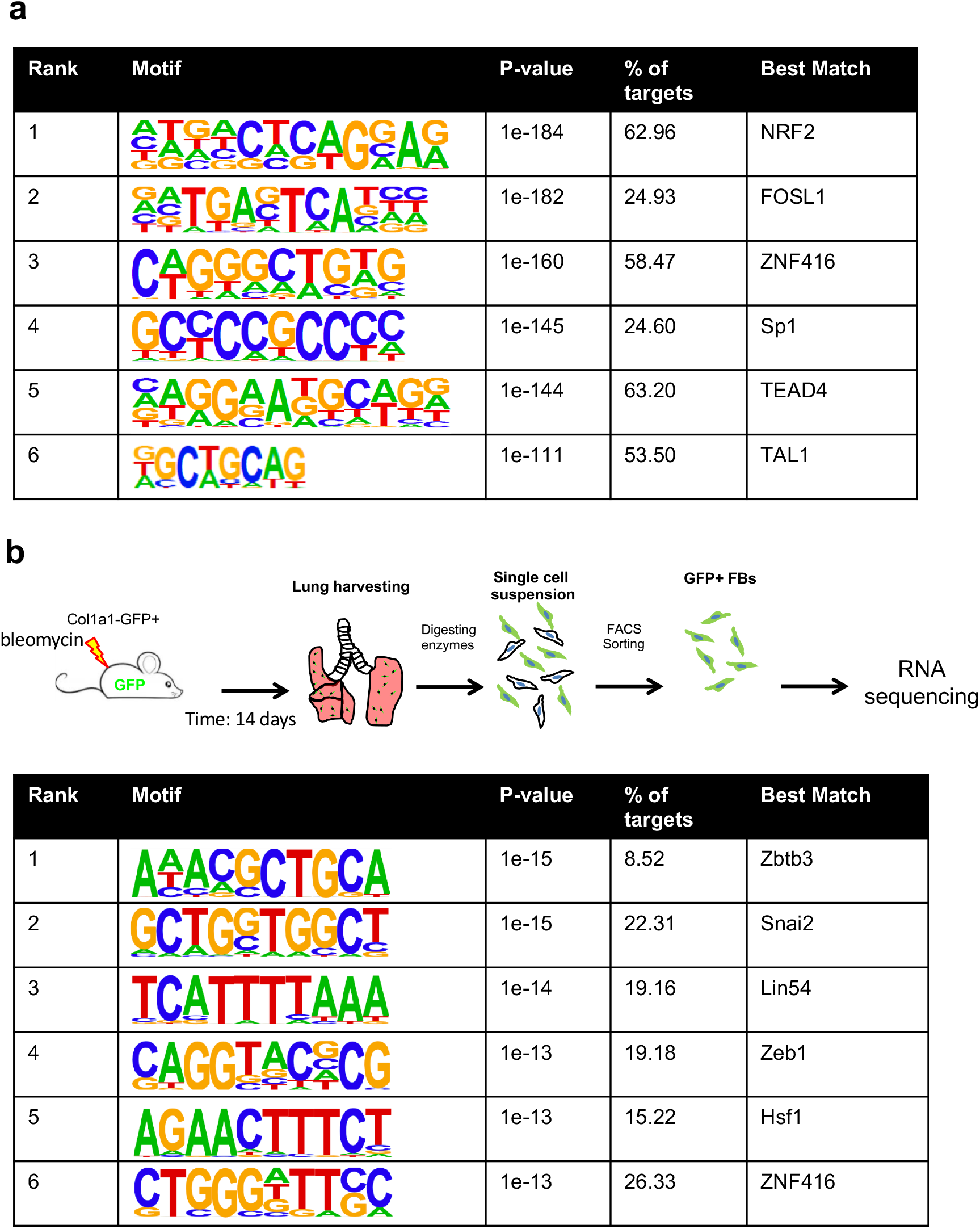
Motif analysis identifies ZNF416 as a putative regulator of fibroblast activation *in vitro* and *in vivo*. (a) *De novo* DNA motif enrichment analysis of differential chromatin accessibility loci of fibroblasts on soft vs stiff matrices ranked by p-value. % of targets indicates the % of total input sequences which contain that respective motif. (b) Schematic of approach to identify differentially expressed genes in mouse fibroblasts following bleomycin challenge to induce experimental lung fibrosis. *De novo* DNA motif enrichment analysis was done in the promoter region (+/−1kb from TSS) of each differentially expressed gene. P-value describes a statistical measure of the enrichment of the DNA motif within the input genomic DNA sequences compared to randomly generated “background” sequences.

To assess the potential roles of these transcriptional regulators in contributing to fibroblast activation *in vivo* we performed motif enrichment analysis scanning promoters (+/−1 kb from TSS) of genes differentially expressed in lung fibroblasts following bleomycin-induced experimental lung fibrosis in mice (Fig. 2b). Interestingly, the top-ranked transcription-factor enriched in both data sets was ZNF416 (Fig. 2b), suggesting ZNF416 may play an unappreciated role in mechanoregulation of fibroblasts *in vitro* and *in vivo*. ZNF416 is a C2H2-type zinc-finger transcription factor and contains a Kruppel-associated box (KRAB) domain and 12 tandem C2H2 Zn finger domains, suggesting ZNF416 can function as both a transcriptional activator and a repressor^46^. The DNA binding motif for ZNF416 was identified in a recent study by Najafabadi, et al^47^. However, beyond its binding motif and protein structure, nothing has previously been reported about ZNF416 biological roles or regulation.

### ZNF416 globally occupies and regulates key genes involved in fibroblast quiescence and activation

To confirm that ZNF416 directly occupies regulatory genomic loci of genes central to fibroblast function, we performed chromatin-immunoprecipitation followed by next-generation sequencing (ChIP-seq) in IMR90 human lung fibroblasts stably expressing ZNF416-FLAG (Fig. S4). ChIP-seq for FLAG identified ZNF416 uniquely binds to 1,386 locations in the human genome. To examine the types of regulatory regions ZNF416 binds to, we annotated ZNF416’s binding sites to the nearest TSS using Homer (Table S6). Annotation revealed ZNF416 predominantly binds to intronic regions (40% of binding sites) and intergenic regions (30% of binding sites), followed by promoter regions (15% of binding sites) (Fig. 3a). Interestingly, ZNF416 largely binds to distal regions of the genome (> 1kb away from the nearest TSS), highlighting a consistency between ZNF416 occupancy and previously reported occupancy of YAP, SMAD3, and SRF (Fig. 3b)^35–37,48^.

**Figure 3:**
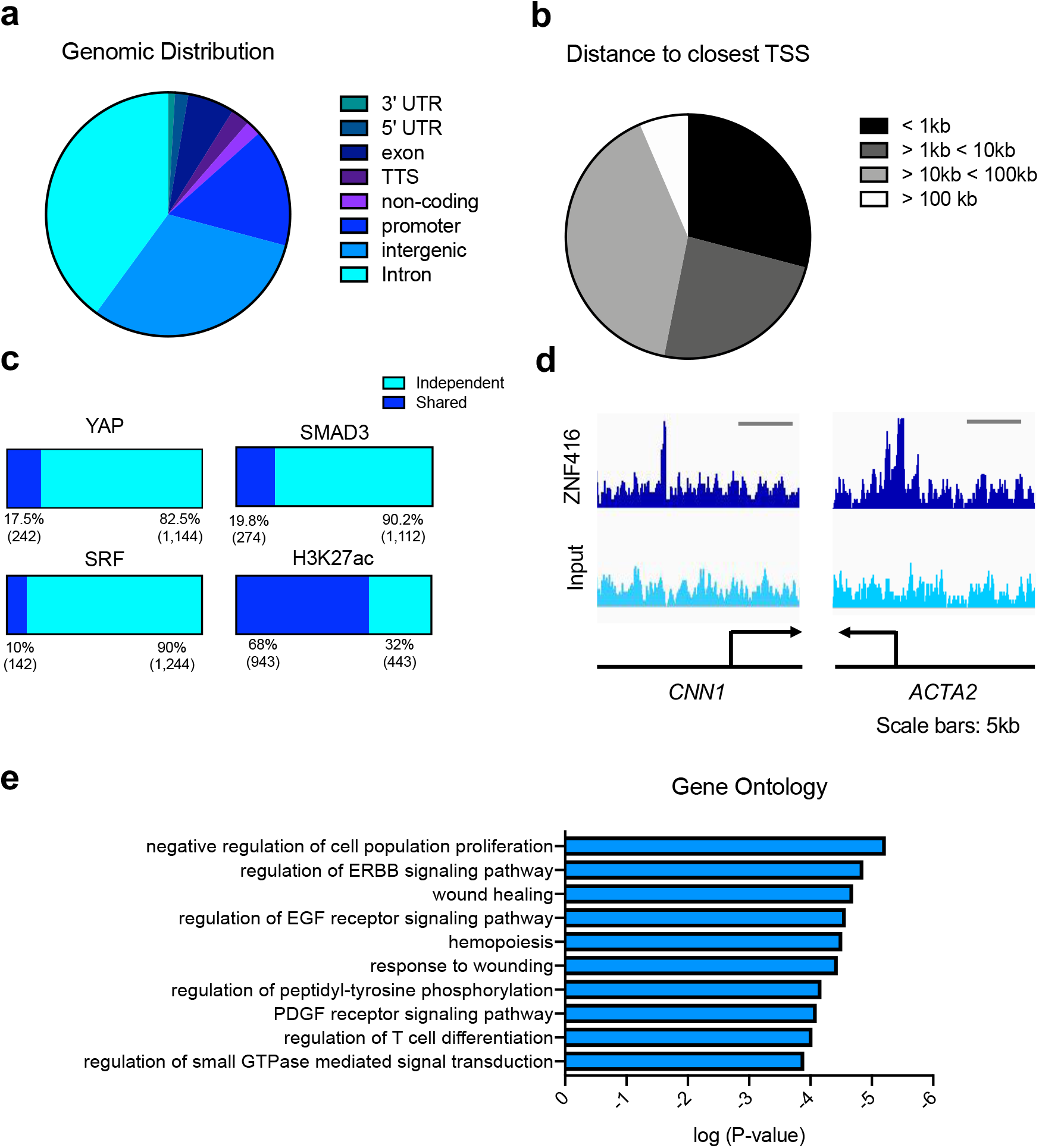
ZNF416 genomic occupancy is distinct from other pro-fibrotic factors and is associated with wound healing and fibrogenic activation programs. (a) Genomic distribution of ZNF416-FLAG binding sites. (b) Genomic linear distance of ZNF416-FLAG binding sites to its nearest TSS. (c) Comparisons of co-occupancy of ZNF416-FLAG with publicly-available YAP, SMAD3, SRF, and H3K27ac ChIP-seq. (d) Genomic view of example ZNF416-FLAG binding proximal to known pro-fibrotic genes. (e) Ontology analysis of genes annotated to ZNF416-FLAG binding sites.

We next sought to determine whether ZNF416 occupancy overlaps with established transcriptional regulators implicated in mechanosignaling and fibrotic pathologies such as YAP, SRF, and SMAD3^2^. To test this, we analyzed publicly-available YAP, SMAD3, and SRF ChIP-seq data (GSE61852, GSE38103, GSE61475 respectively)^35–37^ and evaluated the % of ZNF416 binding sites which reside within 2 nucleosomes (300 bp) of YAP, SRF, and SMAD3. Surprisingly, only 17.5% (242) of ZNF416 binding sites were in close proximity to YAP binding sites, 19.8% to SMAD3, and 10% to SRF suggesting ZNF416 functions in a fashion that is largely independent of established fibrotic transcriptional regulators (Fig 3c). Due to ZNF416 containing both activating (C2H2 Zn) and repressive (KRAB) protein domains, we next cross-referenced our ZNF416-FLAG occupancy with publicly-available H3K27ac ChIP-seq in IMR90 fibroblasts (GSE16256)^38^ to address whether ZNF416 largely acts as a transcriptional activator or repressor. Interestingly, we found that 943 (68%) ZNF416 binding sites were co-occupied by H3K27ac in human fibroblasts (Fig. 3c), suggesting that ZNF416 predominantly acts as a transcriptional activator.

Annotation of ChIP-seq-identified ZNF416 binding sites to the nearest TSS identified ZNF416 to potentially regulate a broad range of genes central to fibroblast contractility (e.g. *ACTA2*, *CNN1*), ECM remodeling (e.g. *LOXL2*, *LOXL4*, *TIMP2*), pro-fibrotic growth factor signaling (e.g. *PDGFA*, *PDGFB*, and *CTGF*), and proliferation (e.g. *CDCA7*) (Fig. 3d, Table S6) (BigWig coverage files will be deposited in GEO). Using Panther, we then conducted an unbiased ontology analysis of the genes nearest ZNF416 binding identified by ChIP-seq. Gene ontology revealed ZNF416 occupies regulatory regions of gene programs involved in wound healing (e.g. *IGF1*, *COL5A1*)^49,50^, and receptor tyrosine kinase pathways previously linked to fibroblast activation and fibrosis, including the epidermal growth factor (EGF) receptor (e.g. *EGFR*)^51,52^ and platelet-derived growth factor (PDGF) receptor (e.g. *PDGFA*, *PDGFB*, *PDGFC*, *PDGFRB*)^53,54^ (Fig. 3e), consistent with a programmatic role for ZNF416 in regulating genes central to fibroblast pathological function.

### ZNF416 is central to fibroblast activation

We next conducted gain and loss-of-function studies to examine the role of ZNF416 in lung fibroblast function. RNAi-mediated knockdown of ZNF416 in fibroblasts cultured on rigid tissue culture plastic attenuated baseline transcript levels of ECM-related genes (e.g. *COL1A1*, *FN1*), contractility related genes (e.g. *ACTA2*), and also proliferation associated genes (e.g. *MKI67* and *CCNA2*) suggesting ZNF416 plays a pleiotropic role in regulating fibroblast functions associated with fibrotic pathologies (Fig. 4a). To examine whether ZNF416 modulates fibroblast contractile function, we performed gel compaction assays using collagen gels and found ZNF416 knockdown prevented TGFβ-mediated fibroblast contraction (Fig. 4b). Additionally, we examined ZNF416’s role in fibroblast proliferation capacity. Using Ki67 immunofluorescence and cell counting as readouts, we found that knockdown of ZNF416 attenuated proliferation when compared to a non-targeting siRNA control indicating ZNF416 is critical for proliferation capacity (Fig. 4c, Fig. S6).

**Figure 4:**
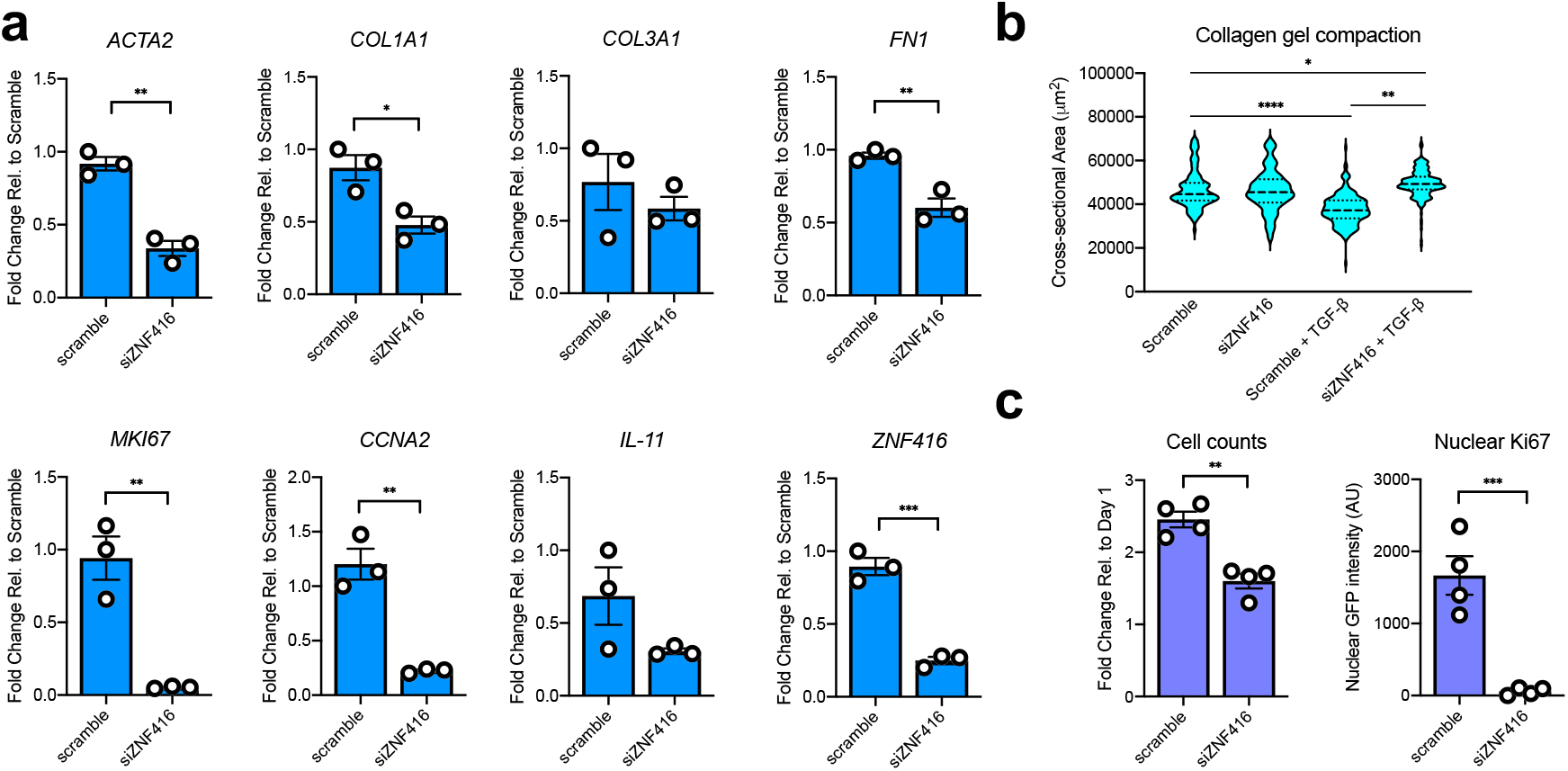
Knockdown of ZNF416 attenuates lung fibroblast activation. (a) qRT-PCR of primary human lung fibroblasts treated with siRNA targeting ZNF416 for 72 hours. n=3 independent biological replicates. (b) Violin plot of cross-sectional area of collagen gels containing fibroblasts with siRNA-mediated knockdown of ZNF416 in the presence or absence of TGFβ. Solid lines represent median. Dotted lines represent upper and lower quartiles. (c) Fold-change in cell counts of primary human lung fibroblasts between 1 and 3 days in culture following knockdown of ZNF416. Data normalized to the cell counts at day 1 to observe proliferation. Ki67 nuclear intensity, assessed by quantitative immunofluorescence, in primary human lung fibroblasts following ZNF416 knockdown. n=4 biological replicates. Data normalized to cell counts per image. *P<0.05, **P<0.01, ***P<0.001, ****P<0.0001 evaluated by unpaired t-test (a and c) or one-way ANOVA (b). Error bars represent S.E.M.

To study gain-of-function of ZNF416, we utilized a lentiviral approach to generate stable over-expression of ZNF416 in primary human lung fibroblasts (Fig. S3). Consistent with RNAi-mediated knockdown of ZNF416, gain-of-function of ZNF416 amplified expression of extracellular matrix-(e.g. *COL1A1*, and *COL3A1*), proliferation associated (e.g. *MKI67*, *CCNA2*), soluble pro-fibrotic signaling (*IL11*), and contractile associated genes (e.g. *ACTA2*) (Fig. 5a) when compared to empty-vector control fibroblasts. Using collagen gels to assay for fibroblast contractile function, we found that overexpression of ZNF416 enhanced contractile function when compared to empty-vector control fibroblasts (Fig. 5b). Interestingly, overexpression of ZNF416 increased contractile function of fibroblasts to the same level as TGFβ-treated empty-vector control fibroblasts. However, ZNF416 overexpression did not alter baseline TGFβ-signaling as measured by phospho-SMAD2 and total SMAD2/3 levels (Fig. S5). Additionally, we found that overexpression of ZNF416 amplified proliferation capacity when compared to empty-vector controls (Fig. 5c, Fig. S7). To test whether gain-of-function of ZNF416 increased ECM deposition, we used an antibody-based method to probe for collagen I and fibronectin deposition. We found overexpression of ZNF416 promoted enhanced deposition of Collagen I and fibronectin (Fig. 5d). Finally, we also studied cells on soft and stiff matrices to test whether gain-of-function of ZNF416 can override the effect of soft matrices. We found that overexpression of ZNF416 on soft matrices elevated ECM, contractile and proliferative gene expression (e.g. *COL1A1*, *ACTA2*, *MKI67*) to levels close to or higher than empty-vector control fibroblasts placed on stiff matrices, suggesting gain-of-function of ZNF416 overrides mechanosignaling from soft microenvironments (Fig. S8). Taken together, these data identify ZNF416 as a mechanoregulator of fibroblast biology and demonstrate that ZNF416 function is critical to fibroblast activation.

**Figure 5:**
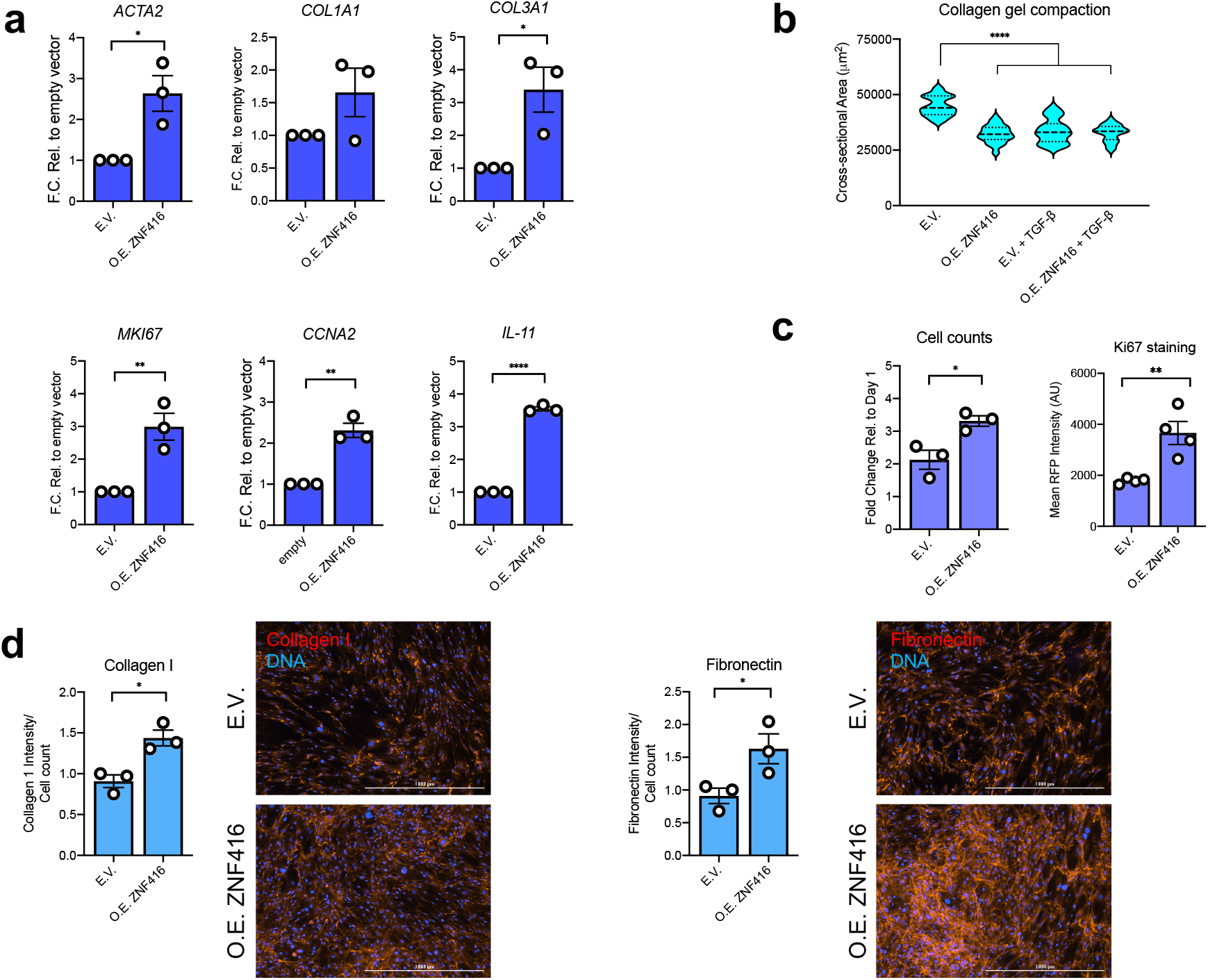
Overexpression of ZNF416 drives lung fibroblast proliferative, contractile and matrix synthetic activation. (a) qPCR of primary human lung fibroblasts stably overexpressing ZNF416 (O.E. ZNF416) or an empty-vector control (E.V.). n=3 independent biological replicates. (b) Violin plot of overexpressing ZNF416 primary human lung fibroblasts and empty-vector controls in the presence or absence of TGFβ. Solid lines represent median. Dotted lines represent upper and lower quartiles. (c) Fold-change in cell counts of primary human lung fibroblasts between 1 and 3 days following overexpressing ZNF416 or empty-vector control. Data normalized to the cell counts at day 1 to observe proliferation. n=4 biological replicates. (d) Deposition of Collagen I and Fibronectin by primary human lung fibroblasts overexpressing ZNF416 or empty-vector control following 3 days of culture. Data normalized to cell counts per image. *P<0.05, **P<0.01, ***P<0.001, ****P<0.0001 evaluated by unpaired t-test (a and c) or one-way ANOVA (b). Error bars represent S.E.M.

## Discussion

Matrix stiffness has emerged as a central modulator of fibroblast function both *in vitro* and *in vivo*, but the transcriptional and epigenetic mechanisms by which matrix stiffness influences fibroblast function remain incompletely understood. Here we demonstrate that *ex vivo* exposure to increased matrix stiffness broadly increases chromatin accessibility in freshly-isolated mouse lung fibroblasts. We identified several transcriptional regulators governing this epigenomic change, including some classified previously as mechanoresponsive (e.g. TEAD4, SP1)^15,55^ and others that are novel, including ZNF416. We confirmed the potential relevance of ZNF416 in an *in vivo* setting through motif analysis of fibroblast genes modulated in the setting of bleomycin-induced lung fibrosis. Investigation of global occupancy of ZNF416 via ChIP-seq in primary human lung fibroblasts revealed ZNF416 targets a wide-range of pro-fibrotic genes including soluble pro-fibrotic growth factors (e.g. *CTGF*, *PDGFA*, *PDGFB*), ECM remodeling proteins (e.g. *LOXL2*, *LOXL4*), and contractility modulators (e.g. *ACTA2*, *CNN1*). Finally, through loss- and gain-of-function studies, we show the importance of ZNF416 in lung fibroblast functions relevant to fibrotic pathologies, including proliferation, ECM deposition and contractility. Taken together, these findings identify ZNF416 as a novel mechano-activated transcription factor that plays a pivotal role in regulating fibroblast functions central to wound healing and tissue fibrosis.

One notable aspect of our work is the use of freshly-isolated fibroblasts to probe the chromatin accessibility changes induced by exposure to matrices of different stiffness. Previous studies that have established transcriptional regulators such as YAP/TAZ and MRTF-A as central mediators of mesenchymal response to mechanical stimuli have used serially-passaged or immortalized cell lines. These proteins have been identified to interact with epigenetic regulating enzymes, such as BRD4, p300, and PCAF^54,56,57^ suggesting these mechanoresponsive transcriptional regulators could mediate lasting mechano-driven epigenomic changes in cells. Indeed, mesenchymal cell populations have been shown to acquire a mechanical memory after prolonged exposure to rigid substrates^16–18,58,59^. Thus, the prior use of serially-passaged cells to examine physiological cell mechanoresponses may have only incompletely identified key factors due to the accumulation of epigenetic mechanical memory in rigid cell culture prior to stiffness control experiments. Our examination of the effect of matrix stiffness on chromatin accessibility of freshly-isolated lung fibroblasts confirmed a role for the YAP/TAZ interacting transcription factor TEAD4, as well as the recently identified matrix stiffness responsive factor Sp1^15^, but also identified ZNF416 as a novel putative mediator of these changes (Fig. 2a). Importantly, ZNF416 was also implicated in our motif analysis of genes regulated *in vivo* in fibroblasts during experimental lung fibrosis, highlighting its potential relevance in wound healing and fibrosis. Moreover, comparison of ChIP-seq data from ZNF416 with other known mechano-responsive and pro-fibrotic transcription factors identified relatively little overlap, suggesting a unique functional role for ZNF416.

ZNF416 contains 12 tandem C2H2 Zn finger domains which have been shown to interact with activating epigenetic regulating enzymes such as p300^60^, a major histone acetyltransferase. ZNF416 additionally contains a KRAB domain, which has been shown to interact with repressive complexes, such as the Nucleosome Remodeling and Deacetylase complex (NuRD)^61^, HP1⍺ (CBX5)^62^, and DNA methyltransferases^63^. The diverse functions of ZNF416’s domains indicate ZNF416 has potential to act either as a transcriptional activator or repressor. However, our cross-reference analysis of ZNF416 and H3K27ac ChIP-seq indicates that ZNF416 largely co-occupies sequences associated with H3K27ac (Fig. 3c), consistent with ZNF416 acting as a transcriptional activator and our observation that gain-of-function of ZNF416 overexpression drives fibroblast gene expression and activation. Taken together, our observations identify ZNF416 as a transcriptional activator that promotes fibroblast proliferative, matrix synthetic and contractile activation.

A key remaining question is whether ZNF416 pioneers alterations in chromatin accessibility, potentially through interactions with epigenetic regulators, or instead is binding to chromatin already opened through independent mechanisms. A recent study observed that mechanical stretch of chromatin was sufficient to induce change in transcription, suggesting mechano-driven changes in chromatin accessibility could be a rapid event^64^. Based on the important functional roles identified here for ZNF416, further elucidation of its roles in epigenetic remodeling and the upstream pathways linking matrix stiffness to its transcriptional functions are likely to provide additional insights relevant to wound healing and tissue fibrosis. In conclusion, our work demonstrates that matrix stiffness is a key driver of increased chromatin accessibility in fibroblasts, and identifies a novel role for ZNF416 in activation of fibroblast gene programs and functions central to pathological fibroblast activation.

## Supplemental Figures and Tables

**Figure S1:**
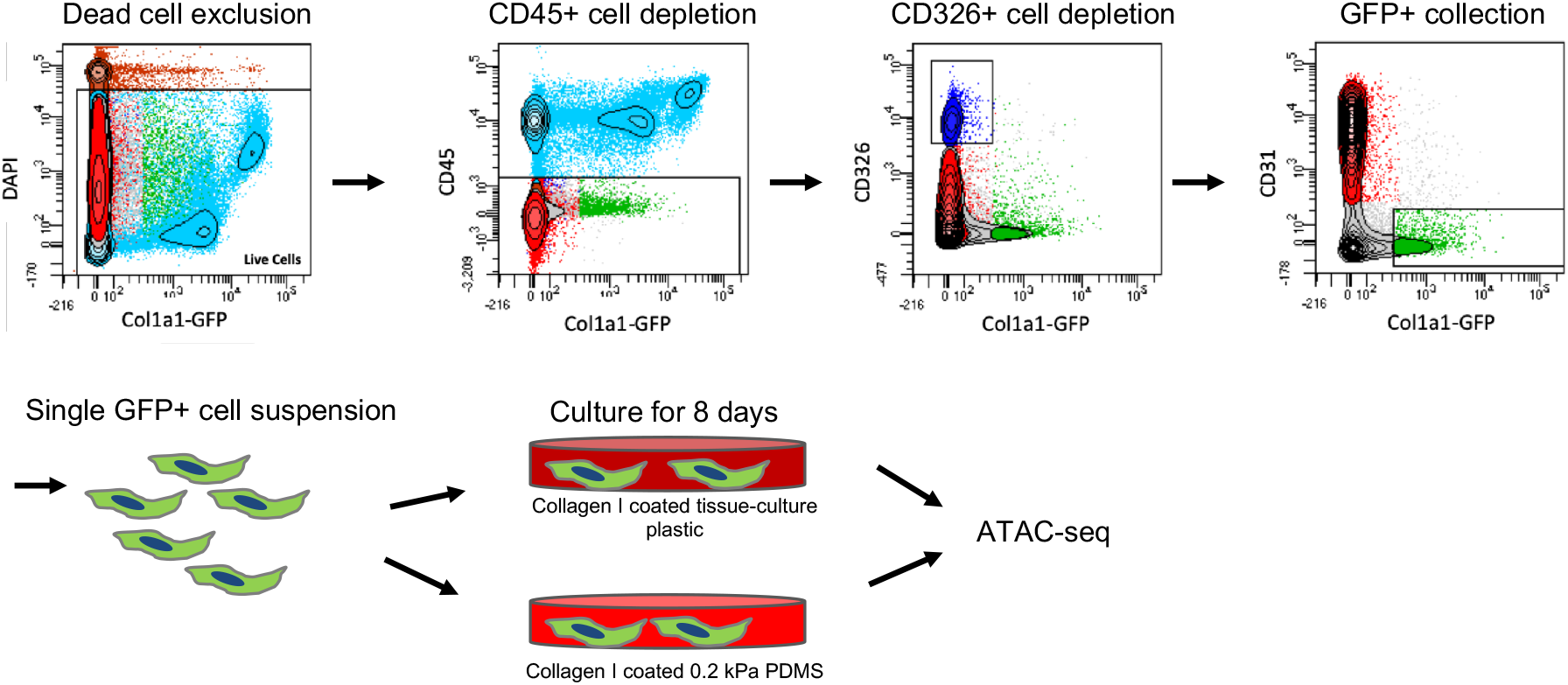
FACS strategy and ATAC-seq approach to examine effect of matrix stiffness on chromatin accessibility of freshly-isolated mouse lung GFP+ fibroblasts. Schematic displaying approach to isolate mouse lung GFP+ fibroblasts. Single cell suspensions were first depleted of dead cells, followed by CD45, CD326, and CD31 depletion, and then followed by a GFP+ selection. Cells were then plated on collagen I coated tissue-culture plastic and 0.2 kPa PDMS for 8 days. Cells were then submitted for ATAC-seq.

**Figure S2:**
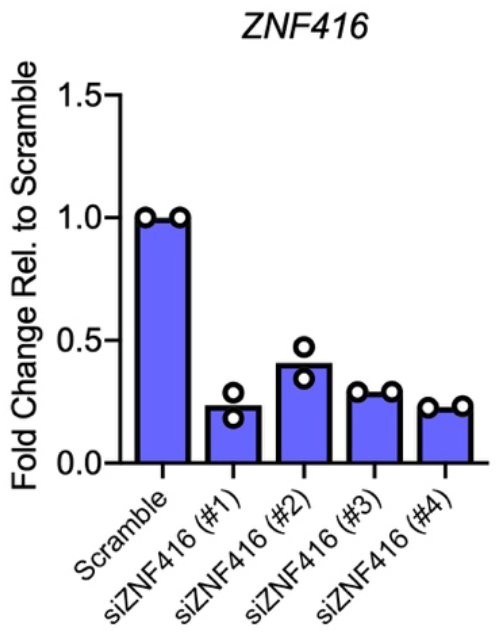
Confirmation of ZNF416 transcript knockdown. qRT-PCR analysis of primary human lung fibroblasts treated with four individual siRNAs targeting *ZNF416* to validate knockdown. Each dot represents an independent biological replicate from a single donor. Data normalized to primary human lung fibroblasts treated with scramble siRNA.

**Figure S3:**
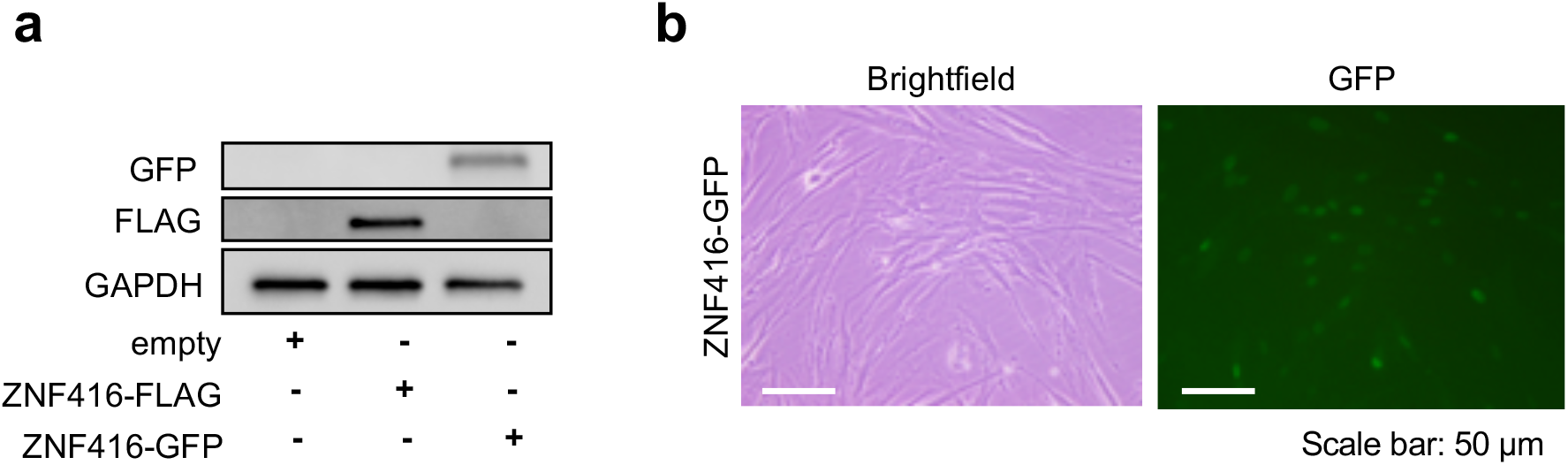
Validation of ZNF416-FLAG and ZNF416-GFP stable expression. (a) Western blot validation for expression of ZNF416-FLAG and ZNF416-GFP in primary human lung fibroblasts. (b) Brightfield and GFP images (10x) of ZNF416-GFP expressing primary human lung fibroblasts. Scale bar represents 50 μm.

**Figure S4:**
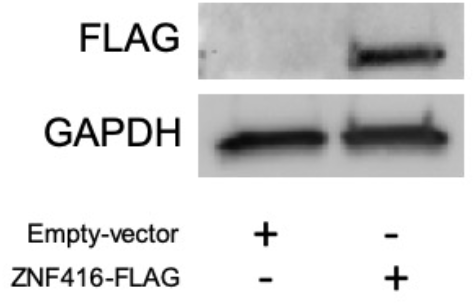
Protein verification of ZNF416-FLAG expression in IMR90 cells. Western blot depicting expression of ZNF416-FLAG in IMR90 cells.

**Figure S5:**
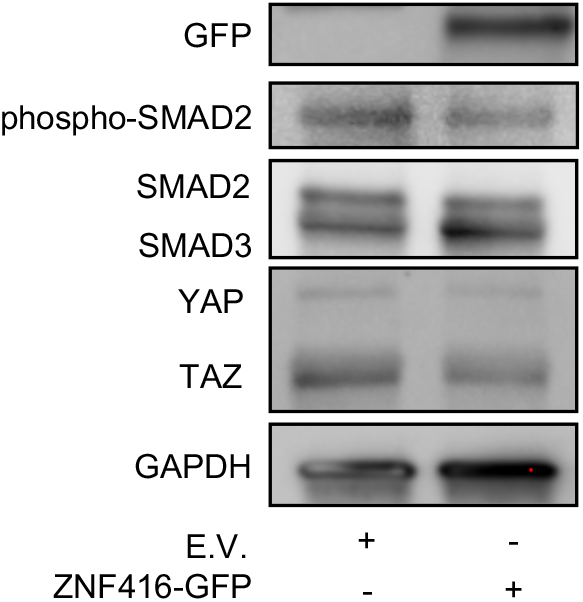
Overexpression of ZNF416 does not alter phospho-SMAD2 and total YAP/TAZ levels. Western blot of empty-vector (E.V.) controls and ZNF416-GFP overexpressing primary human lung fibroblasts.

**Figure S6:**
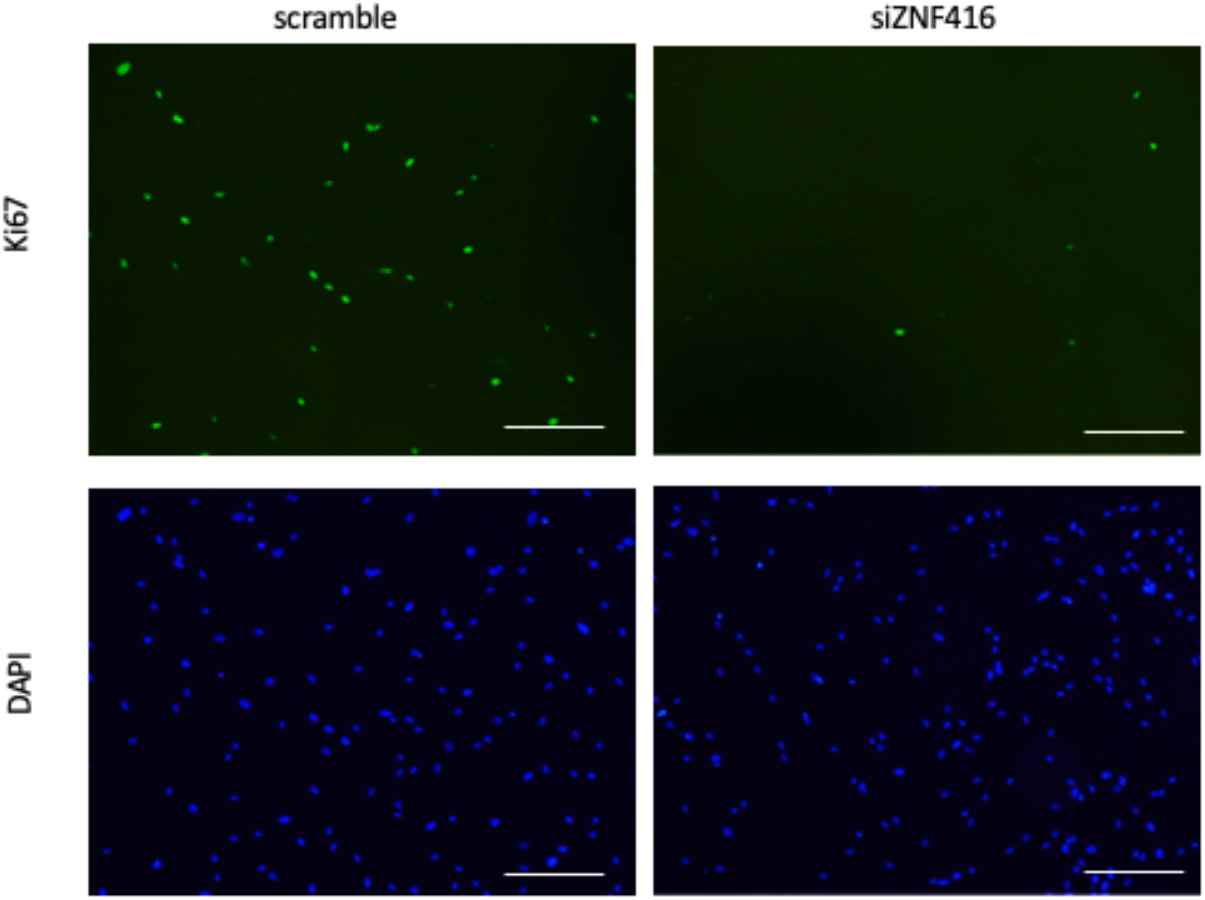
Knockdown of ZNF416 decreases nuclear Ki67 intensity. Representative fluorescence images of Ki67 and DAPI counter-stain of scramble and siZNF416 treated lung fibroblasts. Scale bar is 100 μm.

**Figure S7:**
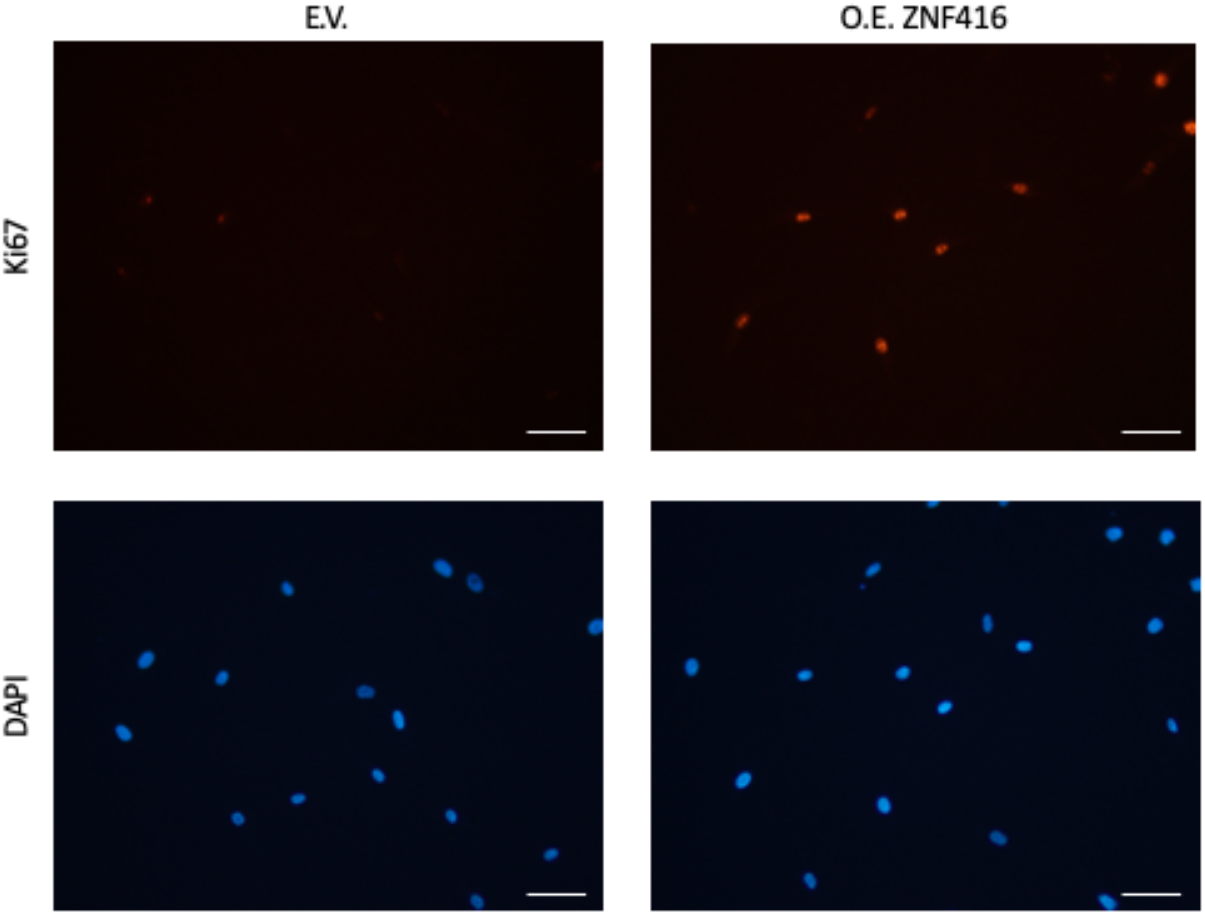
Overexpression of ZNF416 increases nuclear Ki67 intensity. Representative fluorescence images of Ki67 and DAPI counter-stain of empty-vector (E.V.) and overexpressing ZNF416 (O.E. ZNF416) lung fibroblasts. Scale bar is 50 μm.

**Figure S8:**
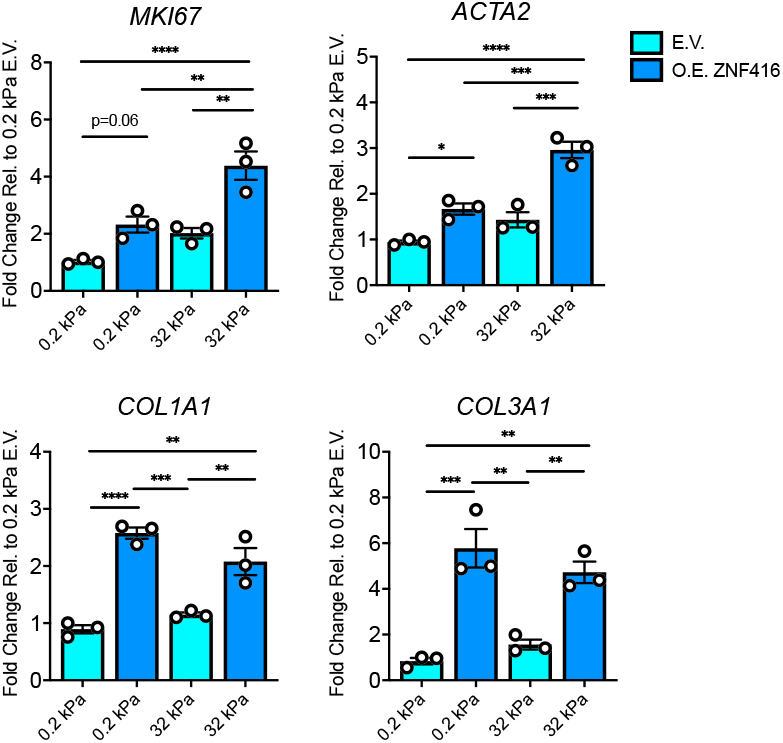
Overexpression of ZNF416 overrides soft matrix effect. qRT-PCR analysis of empty-vector control (E.V.) and ZNF416 overexpressing (O.E. ZNF416) primary human lung fibroblasts placed on collagen I coated soft (0.2 kPa PDMS) and stiff (32 kPa PDMS) matrices for 24 hours. Experiment completed in biological triplicates. *P<0.05, **P<0.01, ***P<0.001, ****P<0.0001 evaluated by one-way ANOVA with Tukey’s correct for multiple comparisons. Error bars represent S.E.M.

