## Supplemental Table 5 for "ZNF416 is a pivotal transcriptional regulator of fibroblast mechano-activation"

| Rank | Motif | P-value | log P-pvalue | % of Targets | % of Background | STD(Bg STD) | Best Match/Details | Motif File |
|---|---|---|---|---|---|---|---|---|
| 1 | 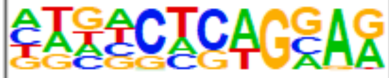 | 1e-184 | -4.248e+02 | 62.96% | 28.71% | 113.0bp (151.9bp) | Nrf2(bZIP)/Lymphoblast-Nrf2-ChIP-Seq(GSE37589)/Homer(0.705) More Information \| Similar Motifs Found | motif file (matrix) |
| 2 | 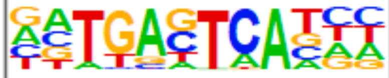 | 1e-182 | -4.196e+02 | 24.93% | 3.68% | 95.4bp (135.0bp) | Fra1(bZIP)/BT549-Fra1-ChIP-Seq(GSE46166)/Homer(0.986) More Information \| Similar Motifs Found | motif file (matrix) |
| 3 | 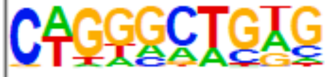 | 1e-160 | -3.684e+02 | 58.47% | 26.90% | 111.7bp (154.0bp) | ZNF416(Zf)/HEK293-ZNF416.GFP-ChIP-Seq(GSE58341)/Homer(0.762) More Information \| Similar Motifs Found | motif file (matrix) |
| 4 | 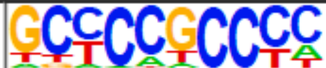 | 1e-145 | -3.358e+02 | 24.60% | 4.87% | 109.6bp (162.0bp) | Sp1(Zf)/Promoter/Homer(0.948) More Information \| Similar Motifs Found | motif file (matrix) |
| 5 | 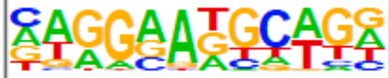 | 1e-144 | -3.331e+02 | 63.20% | 32.60% | 114.8bp (149.9bp) | TEAD4(TEA)/Tropoblast-Tead4-ChIP-Seq(GSE37350)/Homer(0.750) More Information \| Similar Motifs Found | motif file (matrix) |
| 6 | 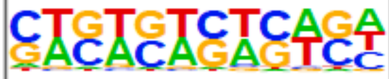 | 1e-130 | -3.002e+02 | 53.55% | 25.46% | 104.6bp (153.3bp) | GAGA-repeat/Arabidopsis-Promoters/Homer(0.749) More Information \| Similar Motifs Found | motif file (matrix) |
| 7 | 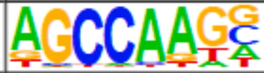 | 1e-111 | -2.575e+02 | 60.50% | 33.62% | 117.2bp (148.5bp) | RIM101/MA0368.1/Jaspar(0.886) More Information \| Similar Motifs Found | motif file (matrix) |
| 8 | 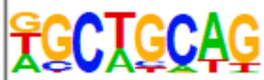 | 1e-111 | -2.567e+02 | 53.50% | 27.34% | 117.8bp (158.8bp) | SCL(bHLH)/HPC7-Scl-ChIP-Seq(GSE13511)/Homer(0.796) More Information \| Similar Motifs Found | motif file (matrix) |
| 9 | 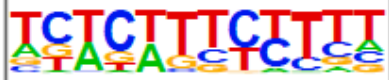 | 1e-95 | -2.205e+02 | 54.64% | 30.13% | 113.0bp (144.2bp) | TEC1/TEC1_YPD/[] (Harbison)/Yeast(0.710) More Information \| Similar Motifs Found | motif file (matrix) |
| 10 | 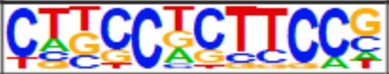 | 1e-73 | -1.699e+02 | 23.89% | 8.60% | 105.1bp (152.2bp) | Gabpa/MA0062.2/Jaspar(0.752) More Information \| Similar Motifs Found | motif file (matrix) |
| 11 | 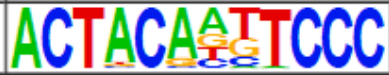 | 1e-71 | -1.657e+02 | 4.92% | 0.03% | 122.3bp (56.3bp) | GFY(?)/Promoter/Homer(0.979) More Information \| Similar Motifs Found | motif file (matrix) |
| 12 | 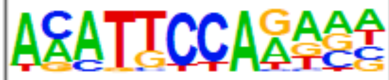 | 1e-66 | -1.542e+02 | 11.87% | 2.34% | 107.3bp (143.3bp) | TEAD(TEA)/Fibroblast-PU.1-ChIP-Seq(Unpublished)/Homer(0.830) More Information \| Similar Motifs Found | motif file (matrix) |
| 13 | 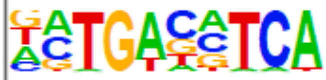 | 1e-62 | -1.443e+02 | 12.91% | 3.04% | 108.2bp (123.1bp) | Atf2(bZIP)/3T3L1-Atf2-ChIP-Seq(GSE56872)/Homer(0.941) More Information \| Similar Motifs Found | motif file (matrix) |
| 14 | 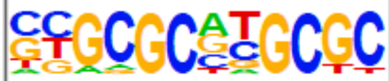 | 1e-54 | -1.253e+02 | 11.83% | 2.92% | 118.0bp (121.9bp) | NRF1(NRF)/MCF7-NRF1-ChIP-Seq(Unpublished)/Homer(0.952) More Information \| Similar Motifs Found | motif file (matrix) |
| 15 | 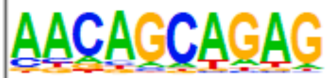 | 1e-52 | -1.206e+02 | 17.41% | 6.23% | 121.1bp (150.2bp) | MyoG(bHLH)/C2C12-MyoG-ChIP-Seq(GSE36024)/Homer(0.703) More Information \| Similar Motifs Found | motif file (matrix) |

| | | | | | | | | |
|---|---|---|---|---|---|---|---|---|
| 16 | 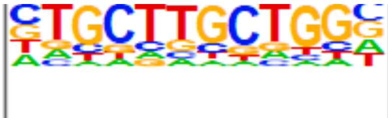 | 1e-43 | -1.000e+02 | 26.06% | 13.17% | 113.7bp (156.0bp) | MBNL1(Znf)/Homo_sapiens-RNCMPT00038-PBM/HughesRNA(0.755) More Information \| Similar Motifs Found | motif file (matrix) |
| 17 | 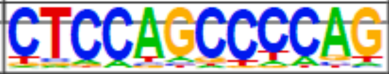 | 1e-39 | -9.011e+01 | 6.76% | 1.27% | 103.0bp (141.9bp) | CRZ1(MacIsaac)/Yeast(0.672) More Information \| Similar Motifs Found | motif file (matrix) |
| 18 | 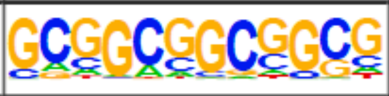 | 1e-36 | -8.358e+01 | 5.77% | 0.97% | 111.3bp (120.6bp) | ERF73(AP2EREBP)/col-ERF73-DAP-Seq(GSE60143)/Homer(0.809) More Information \| Similar Motifs Found | motif file (matrix) |
| 19 | 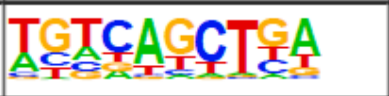 | 1e-35 | -8.181e+01 | 7.95% | 2.00% | 113.5bp (114.0bp) | bZIP52(bZIP)/colamp-bZIP52-DAP-Seq(GSE60143)/Homer(0.795) More Information \| Similar Motifs Found | motif file (matrix) |
| 20 | 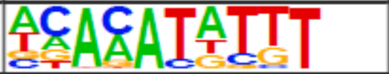 | 1e-29 | -6.826e+01 | 9.18% | 3.06% | 106.8bp (129.2bp) | cad/dmmpmm(Down)/fly(0.818) More Information \| Similar Motifs Found | motif file (matrix) |
| 21 | 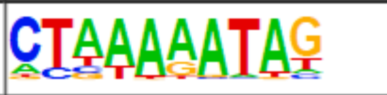 | 1e-29 | -6.694e+01 | 8.66% | 2.79% | 106.6bp (130.6bp) | Mef2c(MADS)/GM12878-Mef2c-ChIP-Seq(GSE32465)/Homer(0.939) More Information \| Similar Motifs Found | motif file (matrix) |
| 22 | 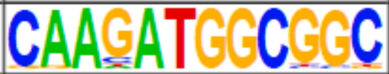 | 1e-27 | -6.359e+01 | 2.32% | 0.09% | 123.8bp (82.0bp) | YY1(Zf)/Promoter/Homer(0.991) More Information \| Similar Motifs Found | motif file (matrix) |
| 23 | 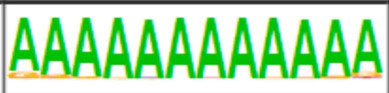 | 1e-27 | -6.330e+01 | 2.13% | 0.04% | 123.5bp (87.5bp) | VRN1(ABI3VP1)/col-VRN1-DAP-Seq(GSE60143)/Homer(0.931) More Information \| Similar Motifs Found | motif file (matrix) |
| 24 | 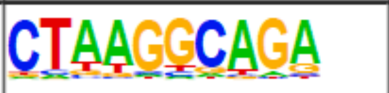 | 1e-23 | -5.372e+01 | 9.13% | 3.55% | 114.1bp (156.5bp) | TBP3(MYBrelated)/col-TBP3-DAP-Seq(GSE60143)/Homer(0.670) More Information \| Similar Motifs Found | motif file (matrix) |
| 25 | 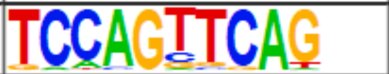 | 1e-20 | -4.637e+01 | 3.64% | 0.73% | 94.6bp (103.8bp) | RORA/MA0071.1/Jaspar(0.678) More Information \| Similar Motifs Found | motif file (matrix) |
| 26 | 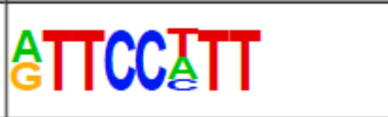 | 1e-15 | -3.519e+01 | 5.82% | 2.23% | 113.1bp (118.8bp) | MOD(RRM)/Drosophila_melanogaster-RNCMPT00140-PBM/HughesRNA(0.839) More Information \| Similar Motifs Found | motif file (matrix) |
| 27 | 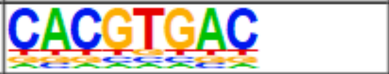 | 1e-14 | -3.228e+01 | 2.46% | 0.49% | 105.7bp (102.9bp) | CBF1(MacIsaac)/Yeast(0.998) More Information \| Similar Motifs Found | motif file (matrix) |
| 28 | 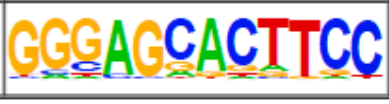 | 1e-13 | -3.019e+01 | 1.94% | 0.31% | 93.9bp (80.9bp) | Znf263(Zf)/K562-Znf263-ChIP-Seq(GSE31477)/Homer(0.740) More Information \| Similar Motifs Found | motif file (matrix) |
